# Directional Variant Tension (T_V_): A Causal Framework for Quantifying Substitution Asymmetry

**DOI:** 10.64898/2026.03.10.710752

**Authors:** Alper Karagöl, Taner Karagöl

**Author notes:** These authors contribute equally. To whom the correspondence should be addressed. Alper Karagöl, Taner Karagöl.

## Abstract

Amino acid substitutions are often directionally asymmetric due to underlying biophysical constraints and diverse evolutionary pressures. We introduce *T*_*ν*_ (variant tension), a kernel regression-based metric that quantifies this directional asymmetry directly from aligned multiple sequence alignments (MSAs). *T*_*ν*_ leverages empirical amino acid frequencies and a non-parametric aussian kernel to capture nonlinear substitution flows, providing a causality-inspired framework for understanding evolutionary dynamics. We also present a web-based application that implements the calculation, allowing users to input MSAs, adjust parameters (kernel bandwidth σ, smoothing window size w), and visualize results, including global tension scores and high-tension sites. Applying *T*_*ν*_ to the human glutamate transporter (EAA1), we identify significant substitution asymmetries, localize high-tension sites, and reveal correlations between elevated *T*_*ν*_ and known pathogenic variants. This framework integrates statistical learning with protein evolution, offering a powerful tool for bridging protein design principles with evolutionary inference. Beyond variant prioritization, T□ offers a scalable framework for simulating evolution under directional constraints, enabling predictive modeling of protein adaptation. The free web application is openly accsessible at https://www.karagolresearch.com/variantt

## Introduction

Understanding co-evolutionary dependencies and directional asymmetries in amino acid substitution is fundamental to deciphering the molecular evolution of proteins, particularly those operating under complex and nonlinear environmental constraints [1,2,3,4,5,6,7,8,9,10]. Historically, the field of molecular phylogenetics has relied heavily on time-reversible, symmetric Markovian processes to model amino acid substitutions - exemplified by classic empirical matrices such as PAM, JTT, and WAG [4,11,12,13,14]. While computationally tractable, these traditional models inherently assume symmetry, thereby overlooking critical directional biases [4]. Biological realities, such as fluctuating selective pressures and shifting structural constraints, frequently drive non-reversible, directional evolution that standard models fail to capture [1,2,3,4,5,6].

Recent efforts to model these asymmetries have explored context-dependent evolutionary models and mutation-selection frameworks; however, these approaches often impose rigid distributional assumptions or suffer from computational overparameterization [4,9,10,11]. In our previous work, we utilized correlation of amino acids in sequence alignments of TLR5s, synaptic proteins, as well as glutamate and monoamine transporters, to detect directional biases [15,16,17,18,19]. Kernel regression is a non-parametric technique used in statistics to estimate the conditional expectation of a random variable [20,21,22,23,24]. Instead of fitting a rigid, predetermined line to the data (like in linear regression), kernel regression finds non-linear patterns by calculating locally weighted averages [20,21,22,23,24]. While conceptually valuable, a major limitation of these early correlation-based approaches was the absence of a robust, standardized metric to quantify and prioritize these differences. The evolution of computational methods for detecting selective pressures has thus reached a critical juncture, where traditional correlation frameworks encounter fundamental limits in both mechanistic interpretation and statistical rigor.

The Directional Variant Tension methodology addresses these limitations through a paradigm shift from inferential statistics to direct probabilistic modeling of amino acid substitution propensities, representing a substantial advancement in phylogenomic analysis capabilities. We define variant tension (*T*_*v*_) as a kernel regression estimator designed to quantify directional flow between residue identities at homologous sites. *T*_*v*_ measures localized substitution tension by comparing nonlinear conditional expectations of amino acid frequencies within a Gaussian kernel smoothing [25,26,27,28] framework. By utilizing a Gaussian function, this framework ensures that closely related evolutionary data points exert an exponentially stronger statistical influence on the estimation than distant ones.

The statistical foundation of the Directional Variant Tension methodology addresses several fundamental issues that compromised the reliability of correlation-based approaches. The inherently non-parametric nature of kernel-based probability estimation eliminates distributional assumptions that frequently violated in amino acid frequency data, providing more robust statistical inference [20,21,22,23,24]. The normalization procedures ensure that tension values remain bounded and comparable across positions and protein families, addressing scale invariance problems that plagued ratio-based metrics in previous methodologies. We demonstrate the utility of this framework by applying it to the human glutamate transporter (EAA1). Our previous studies indicated assymetric substitation in these conserved proteins that may explain diversification of membranous helices [17,29]. Their critical physiological roles and the complex selective pressures acting upon them make them ideal candidates for an in-depth analysis of asymmetric substitution dynamic.

The integration of directional substitution analysis with existing evolutionary genomics workflows promises to enhance our understanding of molecular evolution mechanisms. By providing quantitative measures of substitution asymmetry that complement traditional evolutionary analyses, the *T*□ framework enables more comprehensive characterization of evolutionary pressures shaping protein sequences. This enhanced analytical capability is particularly valuable for understanding evolutionary innovations, functional diversification events, and the molecular basis of adaptive evolution in natural populations.

## Results and Discussions

### Previous simple kernel-causality analysis of TLR5s evolution and the QTY-code

The removal of the transmembrane and intracellular domains in sTLR5 is accompanied by specific changes in the remaining sequence, also showed themselves in the co-evolution data of aligned residues [18]. Kernel-based correlation analysis in this study reveals that the evolutionary relationship between the soluble (sTLR5) and membrane-bound (mTLR5) forms of Toll-like receptor 5 is asymmetric and multidirectional rather than a simple, focused adaptation[18]. The analysis of kernel causality identified specific directional shifts in amino acid composition: methionine (M) and valine (V) show a significant tendency to move from membrane to solvent environments, while histidine (H) and tyrosine (Y) tend to move from solvent to membrane [18]. Furthermore, significant causal directions were established for several substitutions, such as valine leading to threonine (V>T) when regressed on alanine intermediates, phenylalanine leading to tyrosine (F>Y), and arginine leading to lysine (R>K) [18].

The primary advantage of employing kernel-based generalized correlation is its non-parametric nature, which allows it to capture complex, non-linear relationships without relying on the strict distributional assumptions of traditional parametric tests [18,19,20,21,22,23,24]. This is particularly beneficial for evolutionary data that may violate homoscedasticity assumptions or lack a continuous time-series structure. This methodological framework enables researchers to identify which variable acts as the kernel cause, providing a more detailed understanding of evolutionary pathways and selective pressures that simple correlation cannot offer.

While this approach successfully identified statistical associations between amino acid frequencies, it suffered from fundamental interpretive limitations inherent to correlation-based inference. The biological meaning of asymmetric correlation coefficients remains ambiguous, requiring post-hoc mechanistic interpretation that may not reflect underlying evolutionary processes. Furthermore, the treatment of evolutionary conservation as a confounding variable necessitated statistical control procedures that artificially separated conservation effects from the primary evolutionary signal, potentially obscuring biologically meaningful relationships between constraint and substitution directionality.

### Statistical limitations of previous work

The computational architecture of the Directional Variant Tension methodology addresses several practical limitations that have hindered large-scale evolutionary analysis (Figure 1). Previous correlation-based pipelines required integration of multiple external tools including ConSurf homolog identification, MAFFT alignment processing, and R-based statistical analysis packages [15,16,17,18,19]. This multi-software dependency created potential failure points, version compatibility issues, and computational bottlenecks that limited scalability for high-throughput proteomic analysis. The *T*□ framework eliminates these dependencies through a self-contained implementation that processes multiple sequence alignments directly, achieving O(L × n × |A|^2^) computational complexity that scales efficiently with alignment size and sequence number.

**Figure 1:**
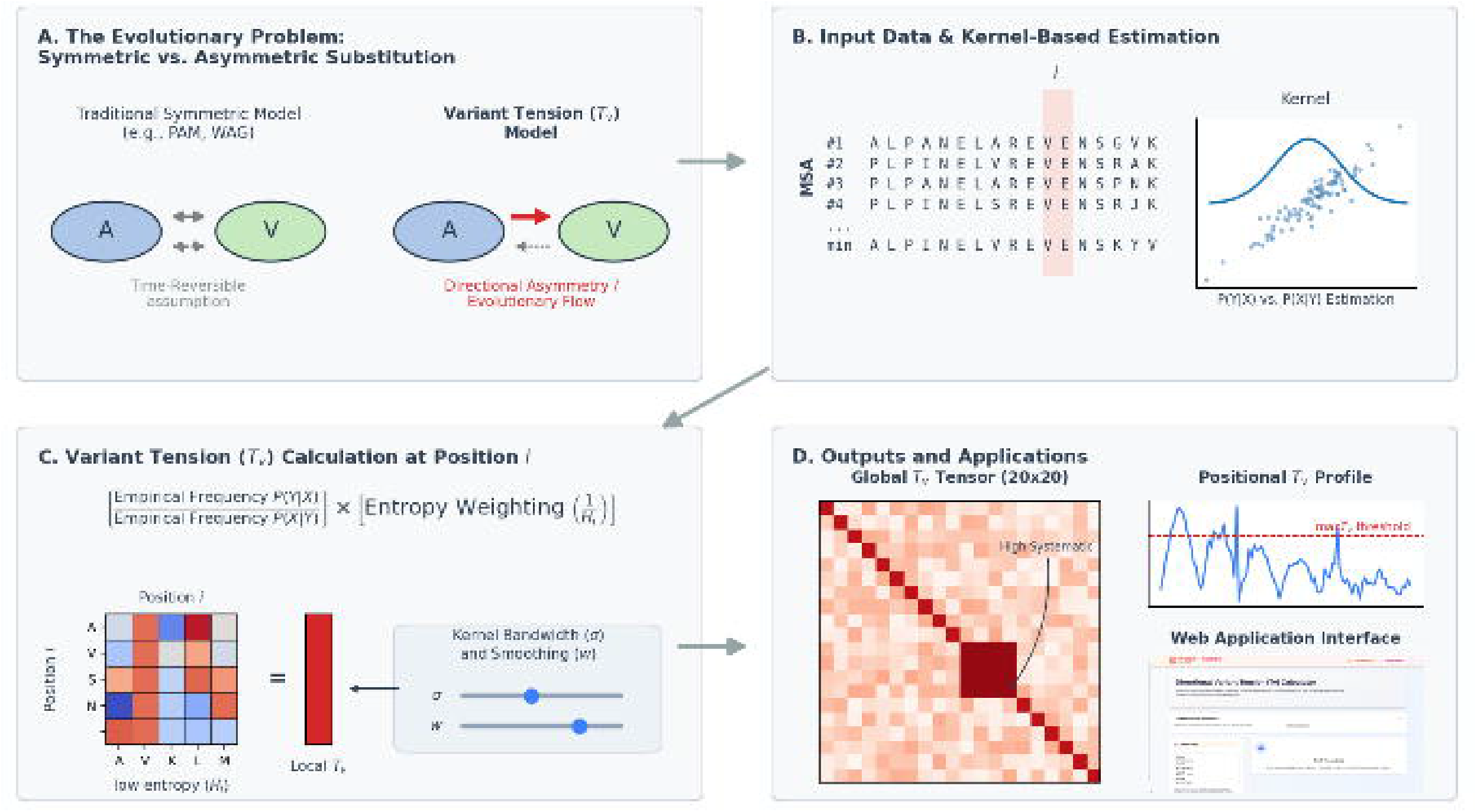
Conceptual Overview and Workflow of the Directional Variant Tension Framework. **\* (A) The Evolutionary Problem:**Contrasts traditional time-reversible, symmetric Markovian models (e.g., PAM, WAG) that often overlook directional biases with the proposed model, which directly quantifies directional asymmetry and substitution flow. **(B) Input Data & Kernel-Based Estimation:** Illustrates the framework’s starting point using a Multiple Sequence Alignment (MSA). It demonstrates the non-parametric Gaussian kernel regression used to estimate the conditional probabilities P(Y|X) and P(X|Y) from empirical amino acid frequencies. **(C) Variant Tension Calculation at Position i:** Shows the computation of localized substitution tension, defined as the ratio of empirical frequencies weighted by the inverse of positional entropy (1/H_i) to prioritize signals at highly conserved sites. This step also highlights the adjustable parameters: kernel bandwidth for sensitivity to frequency variations and smoothing window size for spatial context.**(D) Outputs and Applications:** Displays the primary outputs of the methodology, including the global 20×20 variant tension tensor that maps systematic substitution preferences across the protein. It also shows positional profiling to identify evolutionary hotspots (high-tension sites) exceeding specific thresholds, alongside the interactive web application interface designed for parameter adjustment and visualization.

The kernel-based probability estimation employed in the T□ methodology provides inherent robustness advantages over correlation-based approaches. The Gaussian kernel function K(u,v) = exp(-(u-v)^2^/(2σ^2^)) enables principled handling of sparse amino acid frequency data through controlled smoothing, eliminating the homoscedasticity violations that necessitated non-parametric corrections in correlation-based methods. The kernel bandwidth parameter σ provides transparent control over the bias-variance tradeoff in probability estimation, with clear mathematical interpretation that facilitates systematic optimization across different protein families and evolutionary contexts.

Internal consistency validation represents a significant advancement in analytical rigor. The mathematical relationship R_{Y→X}_ = 1/R_{X→Y}_ provides automatic consistency checking that was absent in correlation-based methods, where asymmetric correlation coefficients could exhibit inconsistent relationships without detection. The integrated MSA quality assessment evaluates sequence diversity, alignment length, and conservation patterns to flag potentially problematic datasets before analysis, preventing spurious results from low-quality input data.

The framework’s capacity for systematic sensitivity analysis addresses a critical gap in previous methodologies, where parameter effects on biological conclusions were rarely evaluated comprehensively. The *T*□ implementation enables real-time parameter exploration and robustness testing, facilitating transparent assessment of result stability across parameter ranges. This capability is essential for establishing confidence in evolutionary inferences and identifying parameter regimes where biological conclusions remain robust despite analytical uncertainties. While correlation-based approaches required empirical optimization of E-value thresholds, sequence identity bounds, and kernel parameters with limited biological meaning, the methodology employs parameters with direct evolutionary interpretation. The kernel bandwidth σ controls the sensitivity to local amino acid frequency variations, while the smoothing window parameter w corresponds to the spatial scale of evolutionary constraints (Figure 2). This transparency enables rational parameter selection based on biological expectations rather than statistical optimization, improving both reproducibility and biological relevance of results.

**Figure 2:**
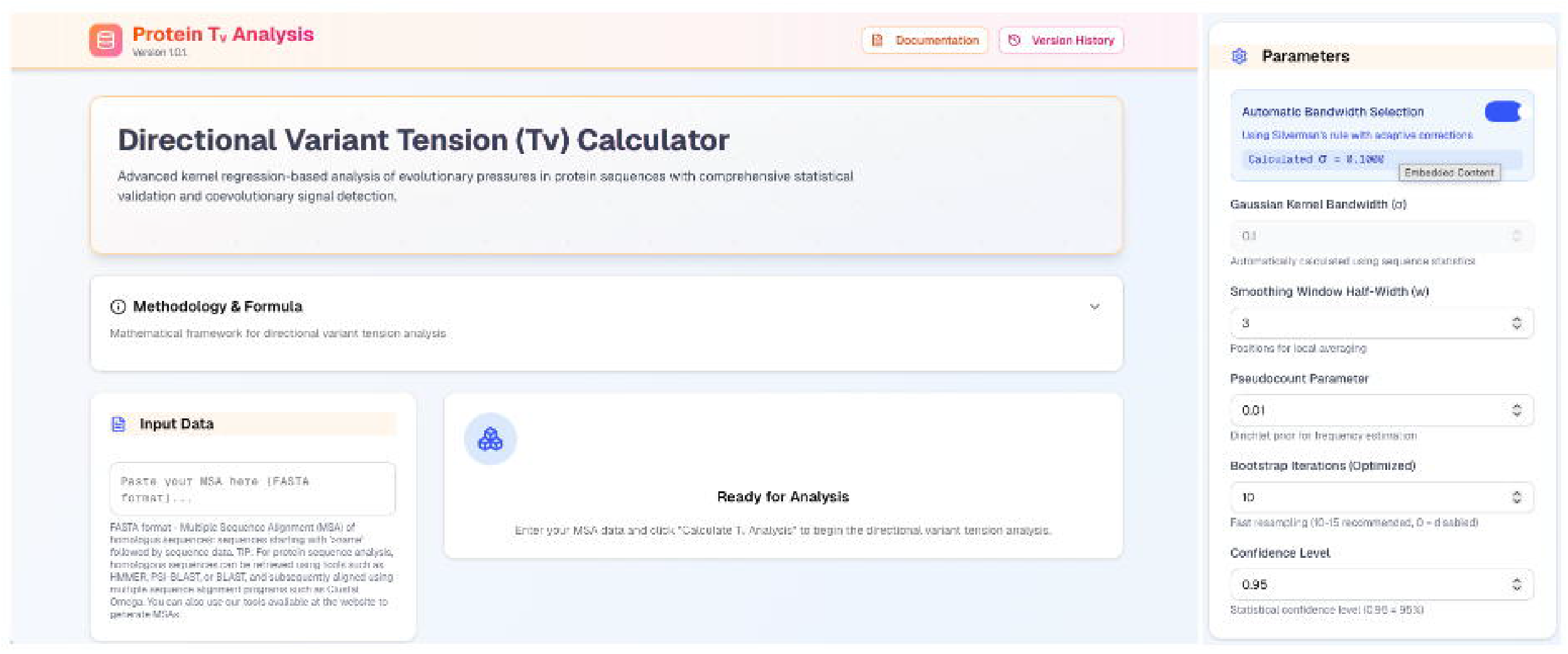
The Directional Variant Tension () Web Application Interface. The web-based calculator, developed using React and JavaScript, provides an accessible platform for executing the kernel regression-based framework. The main panel allows users to directly input Multiple Sequence Alignments (MSAs) in FASTA format. The website enables real-time adjustment of critical evolutionary hyperparameters. Users can manually define or automatically calculate the Gaussian kernel bandwidth (), which controls the estimation’s sensitivity to local amino acid frequency variations

### Mechanistic interpretion

The direct mechanistic interpretation afforded by the Directional Variant Tension framework represents perhaps its most significant advantage over correlation-based approaches. While asymmetric correlations require complex post-hoc interpretation to generate biological hypotheses, tension tensor elements T_{X,Y}_ immediately indicate the relative favorability of specific substitution directions at each alignment position. High tension values directly suggest underlying selective pressures, mutational biases, or structural constraints that favor particular amino acid replacement pathways, providing testable hypotheses for experimental validation.

The multi-scale analytical capability enables detection of evolutionary patterns that remain invisible to position-independent correlation analysis. Regional tension profiles can identify evolutionary domains within protein sequences, revealing functional modules that experience coordinated selective pressures. Global tension patterns across protein families can illuminate universal substitution preferences that reflect fundamental biochemical constraints, while family-specific deviations may indicate adaptive evolutionary pressures unique to particular functional contexts.

The integration of information-theoretic weighting provides biologically meaningful prioritization of evolutionary signals. Positions with low entropy (high conservation) that nevertheless exhibit directional substitution preferences represent particularly interesting evolutionary scenarios, potentially indicating sites where strong functional constraints permit only specific types of amino acid replacements. This treatment of the conservation-variability relationship enables more sophisticated evolutionary hypotheses than simple conservation scoring approaches.

### Directional Asymmetry Captured by Kernel Regression

Our analysis revealed striking directional asymmetries in amino acid substitutions within the transmembrane domains of EAA1 (Table 1, Figure 3). These asymmetries were most prominent in transmembrane helices TM3, TM7, and TM8, regions critical for transporter function 17,29,30,31].

**Table 1.** Glutamate Transporter **1 ( EAA1) High Tension Sites**

| **Position** | **max_tv** | **Direction** | **z_score** | **p_value** |
| --- | --- | --- | --- | --- |
| 429 | 20.6941 | I→C | 1.8954 | 0.058041 |
| 531 | 20.4008 | E→C | 1.7512 | 0.079907 |
| 532 | 20.317 | E→C | 1.71 | 0.087259 |
| 142 | 20.0973 | I→D | 1.6021 | 0.109144 |
| 337 | 20.0966 | L→C | 1.6017 | 0.10922 |
| 336 | 20.0602 | L→C | 1.5838 | 0.113236 |
| 430 | 20.0468 | I→C | 1.5772 | 0.114743 |
| 428 | 20.0422 | I→C | 1.575 | 0.115264 |
| 294 | 19.9363 | L→C | 1.5229 | 0.12778 |
| 533 | 19.8147 | E→C | 1.4631 | 0.143427 |

**Figure 3:**
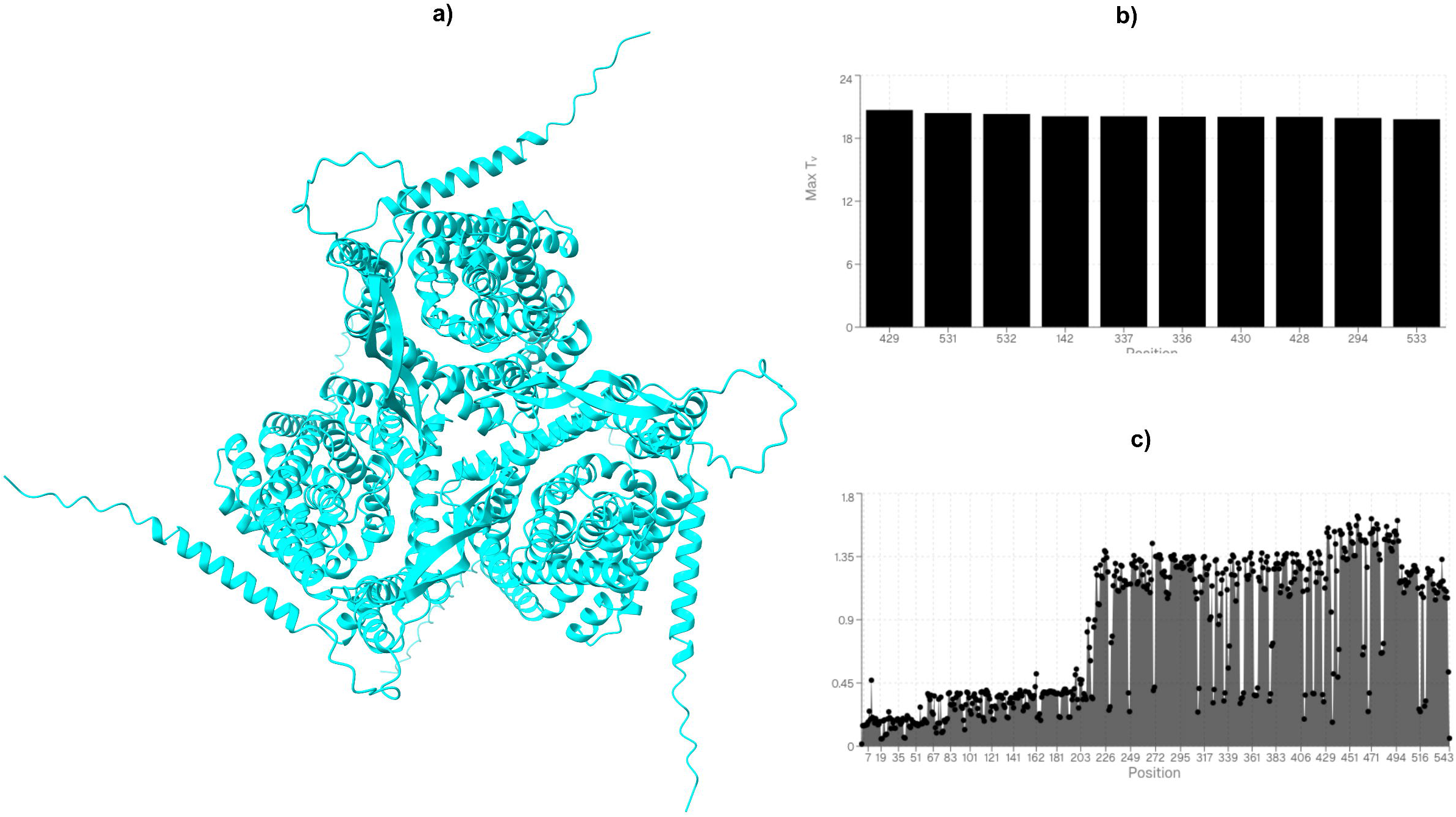
Directional Variant Tension () Analysis of the Human Glutamate Transporter (EAA1/SLC1A3). **(a) Structural Context:** A 3D structural representation of the EAA1 transporter trimer. The framework allows resulting tension patterns to be evaluated directly against known protein structural features. EAA1 represents a structurally conserved neuronal transporter operating under complex selective pressures. **(b) Maximal Tension Analysis:** A ranking of alignment positions exhibiting exceptional directional tension (maxTv). This analysis identifies evolutionary hotspots where specific substitution directions are strongly favored, indicating underlying selective pressures or structural constraints at these specific sites. **(c) Positional Profiling:** A comprehensive positional profile mapped across the sequence length. This multi-scale analytical capability reveals regional profiles that can identify functional modules experiencing coordinated selective pressures, distinguishing between structurally mutable domains and highly constrained microenvironments.

An analysis of high-frequency population polymorphisms reveals a direct concordance with regions of pronounced evolutionary plasticity. The most prevalent missense variant within the population, p.Glu219Asp, exhibits an allele frequency of 1.15% in GnomAD, establishing it as a stable population polymorphism. Thermodynamic evaluation of this specific substitution yields a moderate global value of 4.80, indicative of a chemically conservative transition between isosteric acidic side chains. Positional profiling corroborates this benign biophysical profile, demonstrating that residue 219 is situated within a highly mutable structural domain characterized by an elevated Shannon entropy of 1.51 and an average tolerance threshold (avg*Tv*) of 4.70. Consequently, the protein’s native topological ensemble readily accommodates this biochemical transition without perturbing global thermodynamic stability, perfectly aligning with the variant’s consensus clinical classification as benign.

### Purifying Selection at Structurally Constrained Microenvironments

Conversely, interrogating the ultra-rare variant spectrum elucidates the stringent biophysical thresholds governing critical structural microenvironments. Variants such as p.Leu99His and p.Gly12Trp, existing as mere singletons with extreme minor allele frequencies (approximately 6.84 x 10^-7), introduce severe steric clashes and profound deviations in localized hydrophobicity. The global thermodynamic matrix penalizes these substitutions heavily, assigning them highly disruptive causality scores of 10.36 and 8.92, respectively. When mapped to their specific structural coordinates, positional profiling reveals intense evolutionary constraint; position 99 operates under a restrictive entropy of 0.39, while position 12 exhibits an entropy of 0.43. The superimposition of a high-penalty thermodynamic substitution onto a low-entropy, structurally immutable coordinate invariably induces destabilization of the tertiary fold. As predicted by these metrics, such destabilizing mutations are subjected to intense purifying selection, leading to their near-complete eradication from the human gene pool.

### The Cysteine Exception and Non-Redundant Functional Bottlenecks

The limitations of relying exclusively on global thermodynamic matrices are starkly highlighted by variants operating under strict functional, rather than purely steric, constraints. A prime paradigm of this phenomenon is the p.Cys186Ser mutation, which is definitively classified as a pathogenic variant. From a localized thermodynamic perspective, the metric evaluates the C→S transition as highly conservative, assigning a negligible penalty of 0.82 due to the isosteric nature of the sulfhydryl-to-hydroxyl substitution. However, positional profiling resolves this apparent paradox: residue 186 is governed by a profoundly low evolutionary entropy of 0.37 and a constrained average tolerance (avgTv) of 4.00. This absolute evolutionary intolerance to variation, irrespective of the minimal local steric penalty, strongly implies the residue’s indispensable role in higher-order functional mechanics. The data suggests this cysteine residue likely participates in a non-redundant disulfide bridge essential for maintaining correct conformational architecture or catalytic integrity. Thus, pathogenesis in this instance is driven not by local biophysical perturbation, but by the systemic abrogation of a fold-defining biochemical interaction.

### Implications for Protein Design

The integration of localized evolutionary entropy with global thermodynamic substitution penalties establishes a highly predictive, empirically validated blueprint for rational protein engineering. By leveraging the empirically derived maxTv and avgTv thresholds as absolute topological constraints, protein engineers can rapidly delineate a protein’s viable sequence space. This allows for the precise identification of highly permissive structural domains capable of absorbing radical biochemical modifications - such as the synthetic installation of pH-responsive electrostatic networks, targeted alterations to the isoelectric point, or the incorporation of non-canonical amino acids - without compromising the thermodynamic stability of the native fold.

Furthermore, this multi-parametric approach systematically maps exclusionary zones critical to preserving non-redundant functional mechanics. As exemplified by the strict evolutionary enforcement of critical disulfide networks, identifying low-entropy microenvironments with artificially low thermodynamic substitution penalties prevents the catastrophic abrogation of fold-defining catalytic or structural interactions. Ultimately, utilizing population-validated evolutionary and biophysical constraints transfigures protein optimization from a stochastic empirical process into a deterministic, algorithmic pipeline, thereby accelerating the computational design of hyper-stable, functionally bespoke synthetic isoforms.

### Implications for Evolutionary Genomics

The methodological advances embodied in the Directional Variant Tension framework have broad implications for evolutionary genomics research. The ability to detect and quantify directional and dynamic selective pressures at amino acid resolution enables more precise identification of functionally important sites, improving the accuracy of structure-function relationship predictions [32,33,34,35,36]. The multi-scale analytical capability facilitates comparative evolutionary analysis across protein families, enabling identification of conserved evolutionary mechanisms and adaptive innovations that distinguish related proteins with divergent functions.

The computational efficiency and self-contained architecture of the T□ methodology enable large-scale proteomic analysis that was previously computationally prohibitive. The capacity to process entire proteomes for directional evolutionary signatures opens new research directions in comparative genomics, evolutionary systems biology, and functional annotation of uncharacterized proteins.

## Discussion

Our *T*_*v*_ formulation establishes a novel, statistical mechanics framework for describing directional substitutional forces in molecular evolution. By quantifying variant tension as a dynamic outcome of evolutionary and structural constraints, *T*_*v*_ directly leverages information encoded within empirical amino acid distributions derived from MSAs. Directional variant tension values should be interpreted in the context of: (1) Magnitude: Higher tension values indicate stronger directional preferences. (2) Conservation context: High tension at conserved positions may indicate functional constraints. (3) Structural context: Tension patterns should be evaluated against known protein structural features. (4) Statistical significance: Results should be validated against appropriate null models or randomized controls. The approach assumes that observed amino acid frequencies reflect underlying substitution propensities, which may not hold under strong purifying selection or in the presence of significant back-mutations. Additionally, the kernel-based estimation requires sufficient amino acid diversity at each position for robust probability estimation.

High directional tension values indicate positions experiencing non-neutral evolutionary pressures. These may correspond to functional sites with specific amino acid requirements. The methodology enables several comparative analyses such as inter-family tension pattern comparison to identify conserved evolutionary mechanisms; temporal analysis of tension changes across phylogenetic distances; functional correlation analysis between tension patterns and known protein properties.

A key advantage of *T*_*v*_ over traditional, static substitution models (e.g., BLOSUM or PAM matrices) is its adaptability. *T*_*v*_ adapts locally to the specific sequence variation observed in an MSA, capturing context-dependent evolutionary pressures. Furthermore, its kernel regression structure allows for continuous modeling of substitutional probability flux, thereby avoiding the need to discretize hidden states or assume specific evolutionary pathways. This makes *T*_*v*_ seamlessly integrable with continuous biophysical features and structural analyses. Our findings underscore a new paradigm: amino acid substitution dynamics can reflect both the inherent designability and the evolutionary embeddedness of specific sequence motifs.

## Methods

### Calculating Raw Tv scores using R packages on existing MSA tools

In our previous studies, we employed the ConSurf server, following established protocols [37,38,39,40]. Homologous sequences were identified via HMMER (E-value < 0.0001) using the UniRef90 database, with sequence identity filtered between 35% and 95%. Multiple sequence alignments (MSAs) were generated using the MAFFT algorithm [41]. We subsequently investigated the asymmetric causal relationships between amino acid occurrences within these alignments. Given that our data likely violated homoscedasticity assumptions, we utilized the generalCorr package in R (v4.4.0). This approach leverages kernel regression, a non-parametric method capable of modeling complex non-linearities without rigid distributional requirements [42]. To isolate direct relationships, we calculated partial correlation coefficients (parcor ijk), treating evolutionary conservation as a potential confounding variable. Causal directionality was determined by comparing the absolute values of generalized asymmetric correlation coefficients. (The R Foundation for Statistical Computing, Vienna, Austria), version 4.4.0 (https://www.r-project.org/).

### Statistical Improvements

The Directional Variant Tension framework fundamentally reconceptualizes previous analytical paradigm by directly modeling the conditional probabilities P(Y|X) and P(X|Y) that govern amino acid substitution propensities. This approach transforms the analysis from statistical association detection to mechanistic process modeling, where the ratio R_{X→Y}_ = P(Y|X)/P(X|Y) provides immediate biological interpretation as the relative likelihood of directional substitution. The mathematical elegance of this formulation lies in its direct correspondence to evolutionary mechanisms: values of R_{X→Y}_ >> 1 indicate strong directional preference for X→Y substitutions over the reciprocal Y→X pathway, immediately suggesting underlying functional, structural, or adaptive constraints that favor specific mutational trajectories.

The integration of information theory through positional entropy weighting represents another crucial advancement over correlation-based methods. Rather than treating evolutionary conservation as a statistical confound requiring post-hoc control, the T□ framework incorporates positional entropy H_i_ = -Σ f_{i,a}_ × log(f_{i,a}_) as an intrinsic component of the evolutionary signal. The inverse entropy normalization (1/H_i_) naturally emphasizes positions experiencing strong functional constraints, where directional substitution preferences are most evolutionarily meaningful. This integration eliminates the artificial separation between conservation and directionality that plagued previous methods, recognizing that constraint and substitution asymmetry represent interconnected aspects of the same evolutionary process. The incorporation of spatial context through sliding window smoothing addresses a critical limitation of position-independent correlation analysis. Previous methods analyzed individual residue positions in isolation, potentially missing functionally relevant patterns that emerge from local structural or functional constraints.

### Empirical Frequency Estimation

Let 𝒮 = {𝒮^(1)^,…,𝒮^(*n*)^} represent an MSA of *n* sequences over *L* positions (*i* = 1,…,*L*). For any amino acid *X* ∈ 𝒜 (the set of 20 canonical amino acids), the empirical marginal frequency at position *i* is defined as: 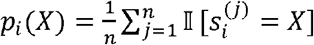 To characterize positional variability, we calculate the position-wise entropy: *H*_*i*_ = –Σ_*X* ∈ 𝒜_ *p*_*i*_(*X*)log(*p*_*i*_(*X*) + *ε*) where *ε* = 10 ^−6^ is a small constant added to prevent logarithmic divergence for zero frequencies.

### Kernel-Based Substitution Tension Estimation

We define a frequency function *f*_*i*_ : 𝒜 → [0,1] that maps each amino acid to its relative frequency at column *i*. For any ordered pair of amino acids (*X,Y*) ∈ 𝒜^2^, the conditional kernel regression estimate of the frequency of *Y* given *X* at position is: 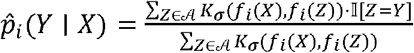 Here, 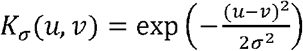 is the Gaussian kernel, where *σ* is the bandwidth parameter.

Directional variant tension at position *i* for the substitution from *X* to *Y* is then defined as: 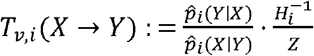 where 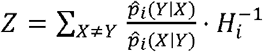 serves as a normalization constant to ensure comparability across positions. This formulation quantifies the relative propensity *X* → *Y* of versus *Y* → *X* substitutions, weighted by the inverse of positional entropy to emphasize highly conserved sites.

### Tensor Representation and Smoothing

For a comprehensive representation of substitution dynamics at each position, we define a variant tension tensor *T*_*v,i*_ ∈ ℝ^20×20^, where *T*_*v,i*_ [*X,Y*] = *T*_*v,i*_(*X* → *Y*). To mitigate noise and capture local positional continuity, we apply a sliding window average: 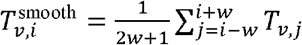 where *w* is the window half-width. At alignment boundaries where the full window cannot be applied, we implement adaptive window sizing:

effective_window_size = min(i+w, L-1) - max(0, i-w) + 1

This ensures consistent smoothing behavior across the entire alignment length while preventing boundary artifacts.

### Global Tension Profiles and Aggregation

The global directional variant tension for a specific substitution *X*→*Y* is obtained by averaging across all alignment positions: 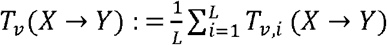 A high *T*_*v*_(*X* → *Y*) indicates a systematic and pronounced substitutional asymmetry favoring *X* → *Y* across the protein, suggesting potential adaptive or selective pressures. Positions exhibiting exceptional directional tension are identified through maximal tension analysis. The corresponding amino acid pair (X*,Y*) represents the most strongly favored substitution direction at position i. Sites are ranked to identify evolutionary hotspots.

### Implementation

A web-based application was developed using React and JavaScript, utilizing Vercel platform. The interface allows MSA input, parameter (σ,w) adjustment, and interactive visualization of global T^−^v and high-tension sites. The kernel bandwidth σ represents a critical hyperparameter balancing estimation bias and variance. Smaller σ values increase sensitivity to local frequency variations but may introduce noise, while larger values provide smoother estimates but may obscure genuine asymmetries.

The algorithm exhibits O(L × n × |A|^2^) time complexity for tension tensor computation, where L is alignment length, n is the number of sequences, and |A| = 20. The smoothing step adds an additional O(L^2^ × w × |A|^2^) complexity. Memory requirements scale as O(L × |A|^2^) for tensor storage.

### Analysis of EAA1

Sequences of human EAA1 (SLC1A3) and their homologs were retrieved from the UniRef90 database. Multiple sequence alignments were generated using Clustal Omega (https://www.ebi.ac.uk/jdispatcher/msa/clustalo) [43]. The alignment in FASTA format converted by Seqret. Missense variants for SLC1A3 were retrieved from gnomAD [44,45]. These variants were mapped to corresponding MSA columns. Each variant was annotated with its local and smoothed *T*_*v*_ value calculated via web application using default parameters. The analysis was conducted using the following computational parameters: A Gaussian kernel was employed for the regression analysis. The Gaussian Kernel Bandwidth () was set to 0.1000, determined through an automatic selection process utilizing Silverman’s rule with adaptive corrections based on the specific sequence statistics of the input Multiple Sequence Alignment (MSA).To facilitate local signal detection, a Smoothing Window Half-Width () of 3 was applied, defining the range of positions used for local averaging across the sequence. To account for sparse data and potential sampling biases in the MSA, a Pseudocount Parameter of 0.01 was utilized as a Dirichlet prior for amino acid frequency estimation. The robustness of the scores was assessed through 10 bootstrap iterations, utilizing an optimized fast resampling approach. Statistical significance and validation were calculated at a Confidence Level of 0.95 (95%). The structure of EAA1 is oredicted utilizing AlphaFold server [46].

### Ethics Approval

Ethics approval was not required for this computational study as it did not involve animal subjects, human participants, and identifiable data.

### Consent to participate

Not applicable. This computational study did not involve human participants.

### Consent for publication

Not applicable. This computational study did not involve human participants.

## Availability of data and materials

Each statistical and computational analysis of this study, included with step-by-step instructions where possible, are publicly available to ensure repeatability. For more detailed information on the statistical analyses, input files and detailed outputs, including codes to regenerate analyses, please visit the website: https://github.com/karagol-alper/Tvariant

## Competing financial interests

None.

## Funding

The author(s) received no specific funding for this work.

## Acknowledgements

None.

## Notes

https://github.com/karagol-alper/Tvariant

### Competing Interest Statement

The authors have declared no competing interest.

